# An Ishihara-style test of animal colour vision

**DOI:** 10.1101/382051

**Authors:** Karen L. Cheney, Naomi F. Green, Alexander P. Vibert, Misha Vorobyev, N. Justin Marshall, Daniel C. Osorio, John A. Endler

## Abstract

1. Colour vision mediates ecologically relevant tasks for many animals, such as mate choice, foraging and predator avoidance. However, our understanding of animal colour perception is largely derived from human psychophysics, even though animal visual systems differ from our own. Behavioural tests of non-human animals are required to understand how colour signals are perceived by them.
2. Here we introduce a novel test of colour vision in animals inspired by the Ishihara colour charts, which are widely used to identify human colour deficiencies. These charts consist of dots that vary in colour, brightness and size, and are designed so that a numeral or letter is distinguishable from distractor dots for humans with normal colour vision. In our method, distractor dots have a fixed chromaticity (hue and saturation) but vary in luminance. Animals can be trained to find single target dots that differ from distractor dots in chromaticity. We provide Matlab code for creating these stimuli, which can be modified for use with different animals.
3. We demonstrate the success of this method with triggerfìsh, *Rhinecanthus aculeatus,* and highlight behavioural parameters that can be measured, including success of finding the target dot, time to detect dot and error rate. Triggerfìsh quickly learnt to select target dots that differed from distractors dots regardless of the particular hue or saturation, and proved to use acute colour vision. We measured discrimination thresholds by testing the detection of target colours that were of increasing colour distances (ΔS) from distractor dots in different directions of colour space. At least for some colours, thresholds indicated better discrimination than expected from the Receptor Noise Limited (RNL) model assuming 5% Weber fraction for the long-wavelength cone.
4. This methodology seems to be highly effective because it resembles natural foraging behavior for the triggerfìsh and may well be adaptable to a range of other animals, including mammals, birds, bees and freshwater fish. Other questions may be addressed using this methodology, including luminance thresholds, sensory bias, effects of sensory noise in detection tasks, colour categorization and saliency.

## Introduction

Over recent years, studies of animal colour vision have focused on the identification of physiological mechanisms, including photopigment and photoreceptor spectral sensitivities and neurons coding for opponency mechanisms, and on theoretical models to predict colour discrimination from this information (e.g. Partridge, 1989; Vorobyev & Osorio, 1998; Shapley & Hawken 2002; Porter et al. 2012). Such data and models cannot replace behavioural tests of colour perception, and of the role of colour in animals’ daily lives. Behavioural investigations have tested detection thresholds (Vorobyev, Brandt, Peitsch, Laughlin, & Menzel, 2001; Olsson, Lind, & Kelber, 2015; Champ, Vorobyev, & Marshall, 2016) and higher-order neural processes, such as colour constancy (Olsson, Wilby, & Kelber, 2016; Wilkins, Marshall, Johnsen, & Osorio, 2016), generalization (Baddeley, Osorio, & Jones, 2001; Kitschmann & Neumeyer, 2005; Scholtyssek, Osorio, & Baddeley, 2016) and categorization (e.g. Jones, Osorio, & Baddeley, 2001; Hanley et al., 2017). However, the underlying mechanisms of these processes remain poorly understood, even in primates (Kelber, 2016). In part, this is because behavioral tests of visual processes with non-human animals are challenging and time consuming. Therefore, novel methods for testing of animal colour vision, quickly and with naturalistic behaviour would be very useful for our understanding of ecological and evolutionary processes.

Working with honeybees, von Frisch (1914) conducted the first behavioral demonstration that non-human animals could identify coloured targets, independent of reflectance intensity. Bees trained to receive a food reward from a small glass dish on blue squares, and then selected blue squares from amongst grey squares that varied in brightness. Many subsequent studies have trained animals to a rewarded colour or pattern (Olsson et al., 2015; Champ et al., 2016; Newport et al., 2017), often using operant conditioning and pairwise or multiple choice discrimination tests. In these experiments, subjects learn a specific colour to receive the food reward. To achieve this, memory of the absolute colour is required, as the animal has to recall the colour learnt in a previous test to distinguish it from the more or less similar alternatives. Also, these methods limit the number of colours can be examined within a reasonable time (Goldsmith & Butler, 2003; Olsson et al., 2015; Champ et al., 2016), and are particularly restrictive in animals that are challenging to train. For example, Champ et al. (2016) took four months to train fish to conduct a pairwise discrimination test, and only three out of seven individuals learnt the task well enough to continue to testing. We have also experienced similar difficulties training fish using a paired-choice test methodology. Some animals may also learn the relationship between presented colours in a paired-choice test rather than a specific colour; for example, in Hemmi (1999), wallabies learnt to choose the colour with the highest wavelength, which enabled the testing of multiple colour combinations in the experiment. Other methodologies, including the spontaneous pecking of spots that varied in size and colour (as per Osorio, Miklosi, & Gonda, 1999 with chicks) have been used; however, it is often difficult to disentangle sensory bias and discrimination abilities in such studies.

In this paper, we introduce an oddity from sample method (Zentall & Hogan, 1974) for testing animal colour vision, which is inspired by Ishihara tests used to identify colour vision deficiencies in humans (Ishihara, 1917; Figure 1i). This offers three very significant advantages over most existing methods. First, the task itself does not require memory of the colour, and consequently more closely resembles most methods used to test human colour thresholds, which are based on simultaneous comparisons of adjacent colours (MacAdam, 1942). Second, one can add uninformative variation (noise) in any direction of choice in the animal’s colour space, which can be used to control task difficulty or to investigate neural mechanisms such as opponent channels. Here we add luminance noise to the background colours to confirm that the fish are using chromatic signals. Third, it is easy to collect at least two separate psychometric measures: accuracy (or error rate), and latency (time) to find the target dot, which can be useful for example in evaluating responses to suprathreshold colour differences. In addition, the method makes it easy to test multiple colours in quick succession without retraining, which is a highly efficient experimental design. Finally, we will see that, at least for the triggerfìsh, this task seems to evoke normal foraging behaviour, making it easy to run and giving some confidence that performance is ecologically relevant.

**Figure 1:**
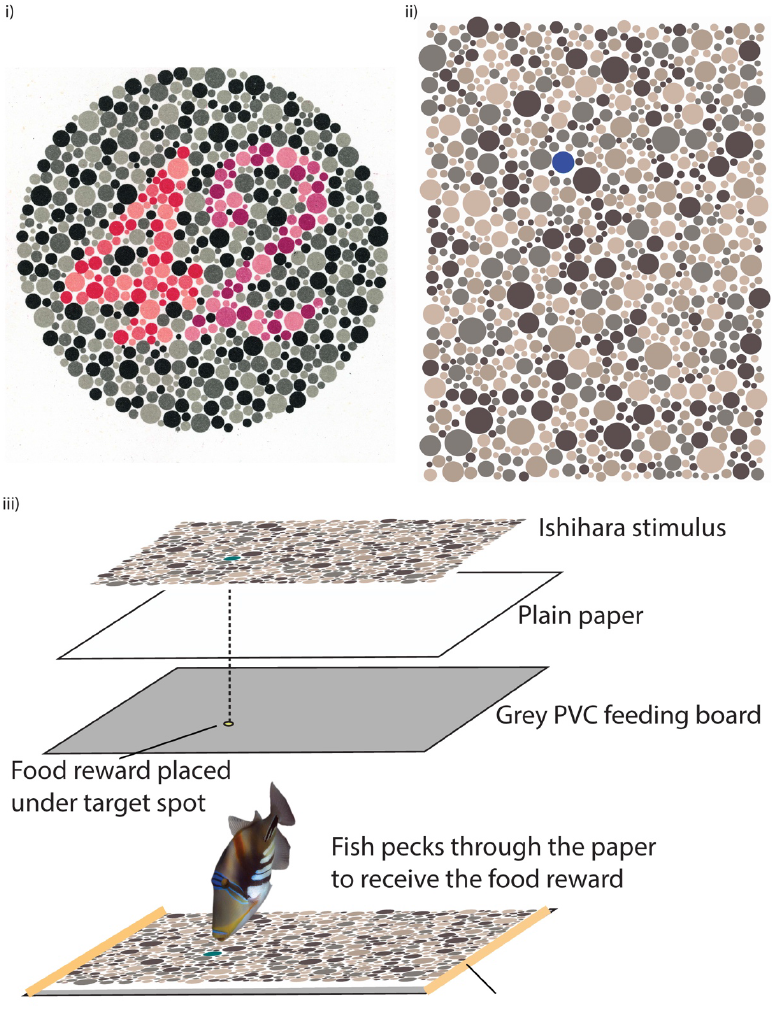
**i)** An example of an Ishihara pseudoisochromatic colour plate (plate 23 of 38 in Ishihara 1917). The background spots vary in luminance and are achromatic, whereas chromatic dots make up the numerical symbols and can be detected due to changes in hue/saturation. Those with normal colour vision will read 42; whereas individuals with strong protanopia (less sensitivity to red light) will read 2, and those with strong deuteranopia (less sensitivity to green light) will read 4. **ii**) An example of our pseudoisochromatic colour stimuli used to test discrimination thresholds in fish. The background dots only vary in luminance are achromatic to triggerfìsh when calibrated and printed on laserjet copy paper. The target dot (in this example, blue) is chosen randomly and varies in hue and/or saturation but is within the luminance range of the background spots, **iii**) Food is placed on a grey feeding board directly under the location of the target dot. A second piece of paper is placed in-between the board and the stimulus to ensure no discernible bump is left by the food that can be detected by the fish. Fish are trained to find and peck through the papers at the target spot to receive the food reward underneath. Elastic bands hold the stimulus in position on the feeding board.

Ishihara plates comprise an array of dots that vary in colour (i.e. chromaticity), brightness and size. The original Ishihara tests of human colour vision exploit the ability of the visual system to segregate elements of an image into figure (object) and ground (background) based on their sharing some common feature, with the colours and brightness of the Ishihara dots being designed so that subjects group dots of (approximately) equal chromaticity, which is dependent on the type of colour vision (Figure 1i). Our tests do not examine visual grouping by colour (although they could easily be modified to do so; Mitchell et al. 2017, Siniscalchi, D’Ingeo, Fornelli, & Quaranta, 2017). Instead the animal learns to find a single target dot that differs in hue or saturation from the background dots that only vary in luminance (Figure 1ii) and tap at it to receive a food reward. We find that the triggerfìsh *Rhinechanthus aculeatus* seems not to learn a particular colour, but instead recognize the odd-one-out (Zentall & Hogan, 1974). This behavior is interesting from the point of view of the fish’s cognitive ecology, and also has the major practical advantage of allowing multiple colours to be tested without retraining fish. We provide a MATLAB code that produces these stimuli and randomizes the location of the target dot.

To demonstrate the performance of the fish in these tests, we provide data to calculate colour thresholds of triggerfìsh and compare these to predictions of the Receptor Noise-Limited Model (RNL model), which is widely used to predict colour discrimination thresholds (just noticeable differences, JND’s) in non-human animals (Vorobyev & Osorio, 1998; Vorobyev et al., 2001).

## Materials and Methods

### Study species

*Rhinecanthus aculeatus* lives on sub-tidal reef flats throughout the Indo-Pacifíc and is a generalist omnivore, feeding predominantly on molluscs and crustaceans. Individuals are easily trained, and perform well in behavioral tests of colour vision (Pignatelli, Champ, Marshall, & Vorobyev, 2010; Cheney, Newport, McClure, & Marshall, 2013; Champ et al., 2016; Simpson, Marshall, & Cheney, 2016; Newport et al., 2017). Fish (n = 8) were collected from shallow reefs around Lizard Island using hand nets, and then shipped to The University of Queensland. Here, they were housed in individual aquaria (60 x 40 x 30 cm deep) with running seawater from a sump and adequate aeration. Tanks were illuminated using KR 96-K36B LED 35W lights (Ecolamps Inc., CA). Experiments were conducted in February-April 2016. Fish were collected under a Queensland General Fisheries Permit (#161624) and a Great Barrier Reef Marine Parks Authority Permit (#G12/35688). This research was conducted in accordance with approval granted by the University of Queensland’s Animal Ethics Committee (SBS/111/14/ARC).

*R. aculeatus* was the first species known to use double cone members independently in colour vision (Pignatelli et al., 2010), and has trichromatic vision based on one type of single cone containing short-wavelength visual pigment (photoreceptor λmax = 413 nm); and a double cone, with one member containing middle-wavelength pigment (photoreceptor λmax = 480 nm) and the other member containing long-wavelength pigment (photoreceptor λmax = 580 nm) (Cheney et al., 2013). *R. aculeatus* has a yellow corneal pigment (Siebeck & Marshall, 2001; Figure S1) whose density increases during the day (Green et al., unpublished data). Because these fish are diurnally active, we therefore modeled the photoreceptor spectral sensitivities with the corneal pigment filtering the incident light. Luminance signals are assumed to be encoded either by both members of the double cone, or by the long wavelength photoreceptor alone (Wild, 2011). In behavioural tests, this species has a visual acuity of 1.75 cycles per degree, similar to goldfish (Champ et al., 2014). In a previous study (Newport et al., 2017), triggerfìsh were able to resolve a pattern of 2 mm (diam.) dots from a control when stimuli were placed from the fish at a similar distance to this study (20 cm or less). The smallest dots in our patterns were 3 mm in diameter, and therefore all spots were visible to the fish at their attack distance (< 20 cm).

### Creating and measuring colours

To calibrate and select our background and target colors, we first created matrices of colours with a range of RGB values using our MATLAB code ‘GetRGBcombinations.m’ (Figure 2). These colours were then printed using a Canon LaserJet Pro 400 printer (Canon, USA) on Steinbeis TrendWhite A4 recycled 80g unbleached white copy paper (Steinbeis Papier GMbH, Glückstadt, Germany). Printing with a LaserJet printer ensured the pigment (actually melted plastic) did not run or change over time when immersed for < 5 min, and no chemicals or dye were released into the water. We chose this paper as it has lower fluorescence than most common brands of bleached printer paper, allowing us to print colours close to the achromatic point, as modelled with the visual system of the triggerfìsh (Figure 3). After printing, the paper was then briefly soaked in water, as the paper would be wet during experiments, removed and the spectral reflectance of each color was measured in air relative to a Spectralon white standard with an Ocean Optics USB2000 spectrophotometer and a desktop computer running OceanView software (Ocean Optics, FL).

**Figure 2:**
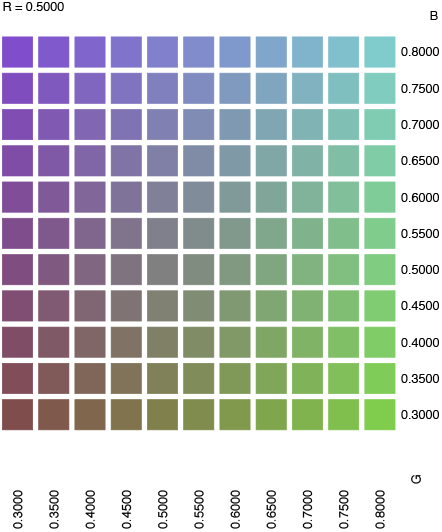
Colour matrix with specific RGB values created using the ‘GetRGBcombinations.m’ code. Colours were printed with a laserjet printer and the spectral reflectance of each colour combination with measured with a spectrophotometer. All the colours in this matrix have an R value of 0.5. The G and B values are shown on the x-axis and y-axis, respectively. We made additional sample colours with other R values ranging from 0 to 1 to make the gamut of experimental colours.

**Figure 3:**
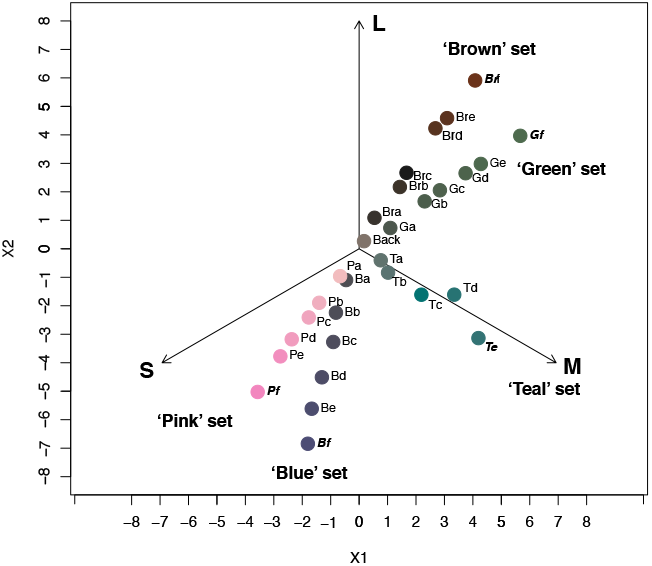
Chromaticity diagram with colour of target dots corresponding to the RNL model: X1 and X2 are defined in equations (4) of Hempel de Ibarra et al. (2001) based on spectral sensitivities of triggerfish *Rhinecanthus aculeatus.* Colours with names in bold-italic *(Brf Gf, Te, Bf, Pf)* were used for training and for reinforcements during testing. Background (Back) denotes the cluster of greys used as the background distractor dots. This plot was produced using the R colour vision package (Gawryszewski, 2017). We will refer to colours approximately on the same line radiating from the achromatic point as colour sets.

Measurements were made using a 200 μm diameter, bifurcated cable, which was also connected to a PX-2 pulsed xenon light source (Ocean Optics, FL). For accurate measurements, the fibre was held 1 mm above the paper at a 45-degree angle with an RPA-SMA Fiber Holder Arm and shielded from stray light. To ensure colours produced by the printer were consistent, each colour was printed on different days and measured on at least five separate occasions (mean difference + st.dev from first printed colour: ΔS = 0.38 + 0.17). Variation was greatest when ink levels of the printer were low, therefore we limited printing of test stimuli to when levels were sufficient. We also used the Colour Calibration and Head Cleaning utility in the printer’s menu regularly to maintain consistency. The Matlab code used to create the colour matrices and pseudoisochromatic stimuli were written by JAE and are available at Dryad Data Repository (http://dx.doi.org/xxxxxx).

### Visual modeling and selection of target colours

Chromaticity of background and target colours was specified by the estimated excitations of triggerfìsh photoreceptors as quantified using photoreceptor spectral sensitivities, illumination and reflectance spectra (Kelber, Vorobyev, & Osorio, 2003). Colour distances ΔS between colours were modelled using the trichromatic photopic receptor noise limited (RNL) model (Vorobyev & Osorio, 1998; Kelber et al. 2003), which assumes that colour discrimination is depended on chromatic signals and limited by noise originating in the receptors. Colours were also plotted in an RNL chromaticity diagram defined in equations (4) (Hempel de Ibarra, Giurfa, & Vorobyev, 2001). We assumed a 1: *2*: 2 ratio (S: M: L) for the weber fraction *(ω)* and the LWS noise threshold was set at 0.05. We use the Weber fraction to estimate noise in the photoreceptors because there are no direct measurements of receptor noise in this species (Kelber et al., 2003). Following evidence from other vertebrates (Vorobyev & Osorio, 1998; Olsson et al., 2015), this model assumes that spatial summation reduces receptor noise, and that noise in each receptor mechanism can be estimated based on the relative abundance of photoreceptor types in the retina. Estimated noise in each photoreceptor was therefore: S 0.07; M 0.05; L 0.05.

Nine reflectance spectra that were very close to the achromatic point were chosen as the background colours (Figure 3, Supplementary Figure S2, S3). Twenty-nine target colours (RGB values used for printing are shown in Table S1) were chosen based on their positions in several radial lines away from the achromatic point (Figure 3); all colours along the same line were similar in hue (line angle from the origin or achromatic point) but varied in chromaticity (distance from achromatic point). On four lines or colour sets (‘Brown’, ‘Green’, ‘Pink’ and ‘Blue’) we used 6 target colours; but on our ‘Teal’ set we used only 5 target colours due to the limitations of laser-jet spot reflectance spectra. All colours were tested in February-April, 2016 with eight fish.

### Experimental setup

During training and testing, tanks were divided into two halves by an opaque partition placed across the tank, which included a door that could be opened by sliding a board upwards. This enabled one side of the tank to be the testing arena, where the stimulus could be set up without fish seeing that area. Printed A4 stimuli were placed on a grey A4-sized plastic (PVC) feeding board and secured in place with two light brown elastic bands at each end, which were ignored by fish during testing. We first trained fish with a plain grey background on which there was only one dot, which was one of the following five colours: *Brf, Gf Te, Bf, Pf* (Figure 3). During the first 2-4 sessions (with 6 trials per session), we placed the food reward (small pieces of squid) on top of the dot to encourage fish to approach and peck at the dot. Once fish began to associate the dot with a reward, the food was then placed on the PVC board, underneath the paper and directly located under the target dot (Figure 1iii). We also placed an additional, plain piece of paper (same stock) under the printed stimuli to ensure that the colour of the stimuli was not altered by the grey PVC board and that no marks or impressions were left by the food. Fish quickly learnt to peck through the paper to create a hole, obtain the food, and spit out the paper. A video of a fish performing this behavior is available in Supplementary Information. Each fish completed a further 5 training sessions with food underneath the dot.

Fish then progressed to a second stage of training during which we used Ishihara style stimuli, which had one target dot that was deemed easily detectable by the fish *(Brf, Gf, Te, Bf, Pf*;, Figure 1ii). Fish learnt the concept that they were to find the dot that differs in hue or saturation (chromaticity) from the background and then peck the target dot to obtain food (squid) placed underneath it (Figure 1iii) within a few days. During this stage, consisting of 12 sessions, fish were randomly presented with the five training colours to ensure they did not learn that the food was rewarded from a particular colour. After 6 training sessions, we found there was no bias in terms of success rate for different training colours (GLMER, z = −1.36, n = 224, P = 0.17).

During testing, trials commenced when the door within the partition was removed, located approximately 30cm from the stimulus, and the fish swam through to the testing arena. Fish were given 30 s to find the target dot and peck through it to receive the food reward (Figure 1iii). Each test session consisted of 5 trials, and 1-2 sessions were conducted per day. The order in which the 29 colours were presented, the size of the dot and position of the target dot were all randomized. In total, we conducted 906 individual trials in March – April 2016 and each target colour was presented to each fish between 2 and 11 times (mean + s.d.: 3.88 + 1.65).

During each trial, we recorded: (1) whether the fish was successful in pecking the target dot within 30 s of entering the testing arena; (2) if so, the time taken from entry to pecking at the target (latency to find the dot); and (3) the number of dots that were pecked incorrectly before the target was pecked. Interestingly, the fish always pecked directly on a dot and not in between dots or elsewhere on the paper. After the target dot had been pecked or 30s had elapsed, the fish were gently encouraged with a net to swim out of the testing arena and back through the door, and the stimulus was removed.

Throughout the experiment, we also randomly conducted 120 control trials (15 per fish) in which there was no differently coloured target dot, i.e. they were all background dots. Food was still placed under one randomly selected dot to ensure that fish were using visual information and not olfactory or other cues other than differences in hue/saturation to detect the target. The mean success rate for control trials was low (3.3%), indicating that, although this was greater than chance (there are approx. 180 of the largest 3 dots on each stimulus and allowing for 5 incorrect choices per 30 s = 0.2%), it was very unlikely they used predominately olfactory or other cues (such as a mark or blister on the paper created by the squid) to find the food.

### Statistical analyses

To model the probability of success for each colour, cumulative Gaussian curves were fitted to the data (Wichmann & Hill, 2001) using the quickpsy package in R (Linares & López-Moliner, 2017). Deviance values were very similar to curves fitted with a logistic curve function. The ΔS at which the probability of success was 50% was calculated for each colour set. In previous studies that use paired choice tests, discrimination thresholds are often modelled at 75% correct choices to be statistically above a 50% random choice threshold (Vorobyev et al., 2001). Due to our experimental design, we reduced our threshold to 50% due to the number of dots being presented to the fish and our control level of 3.3% success rate of detecting the target colour; however, this could be modified depending on the research question. In trials, dot size did not significantly impact the chance of finding the target dot (z = −1.19, n = 906, p = 0.24) as expected.

## Results

For each target colour, the mean success rate at which fish located the target dot ranged from 0% and 100%. For each fish and colour set, the probability of success fitted a normal cumulative distribution function (deviance < 6.31, P > 0.31), with the exception of Fish O-Teal and Fish L-Green, which both exhibited an abrupt step function (Figure 4). Teal had the lowest 50% ΔS threshold (mean + SD: 0.69 + 0.30) and Pink had the highest (2.87 + 0.66). Fish discrimination thresholds for the other colour sets were: Brown 2.33 + 0.35, Blue 2.63 + 0.72, Green 1.39 + 0.57). In successful trials (n = 699), fish took between 1.17 and 29.91 s (mean + SD = 7.04 + 6.43) to find the target dot and fish made between 0 and 8 (0.54 + 1.81) incorrect pecks. During unsuccessful trials (n = 207), fish made between 0 and 7 incorrect pecks (3.05 + 1.91). The average number of incorrect pecks and the time taken to find the target dot decreased with increased ΔS in a non-linear manner (Figure 5). Further analysis of specific threshold data in relation to the fishes’ visual mechanisms is currently being undertaken and will be published in due course.

**Figure 4:**
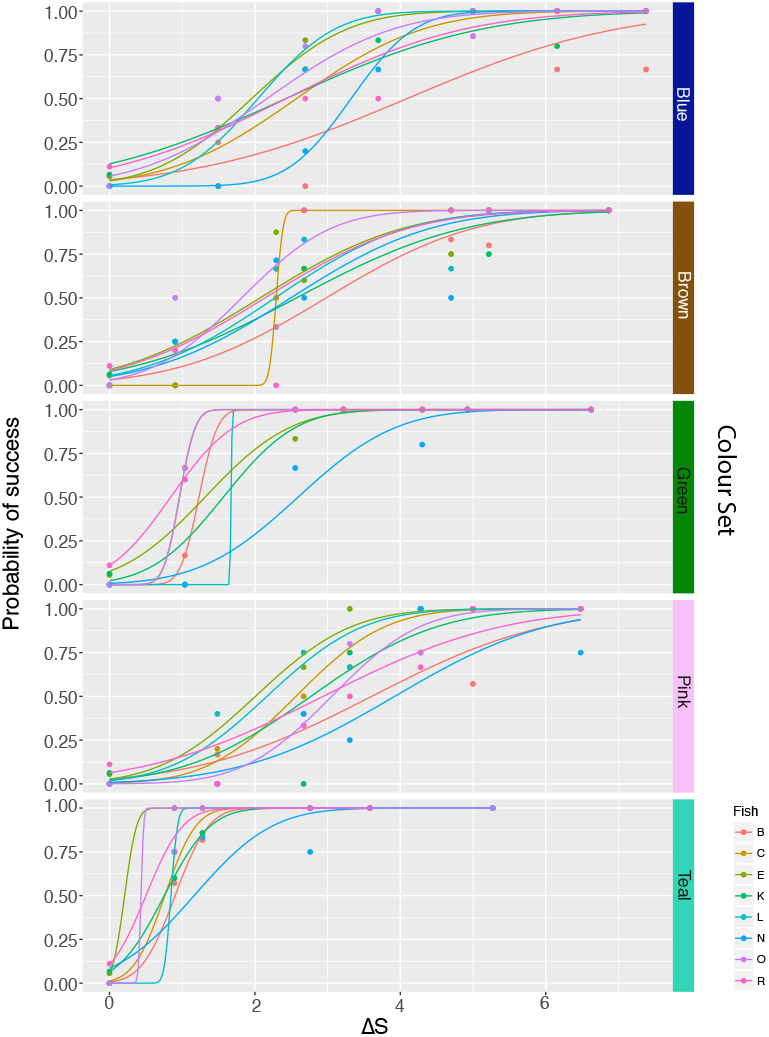
The probability of success in detecting each of the 29 target colours separated by fish and colour set. Psychophysical cumulative Gaussian curves are fitted for each fish with the quickpsy package in R (see text for details). A greyscale version of this figure is available in the supplementary information.

**Figure 5:**
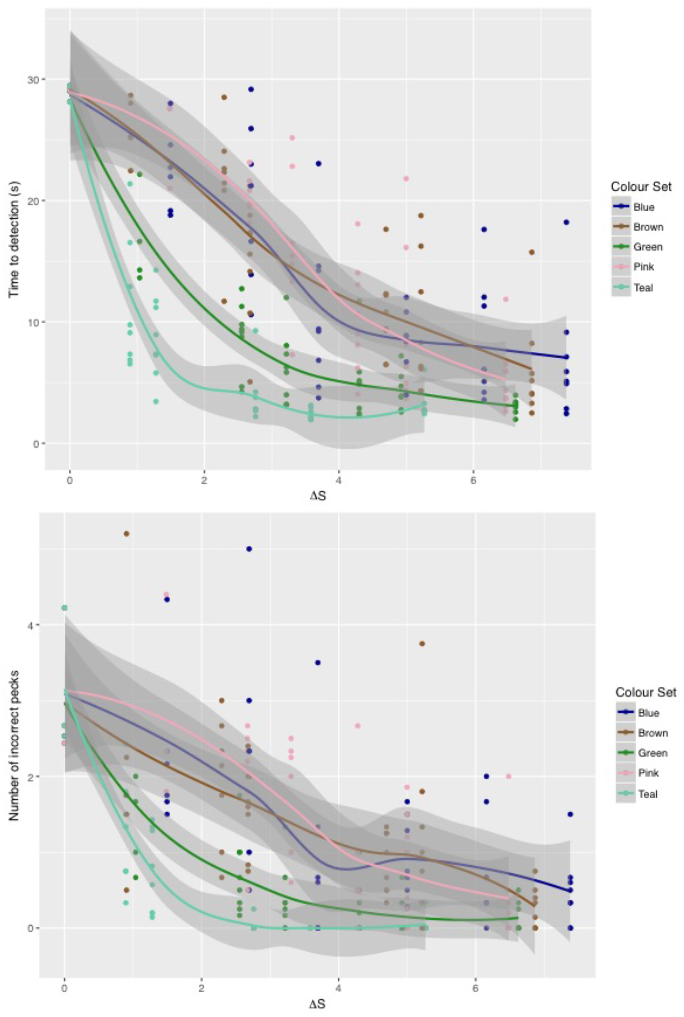
Plots of i) mean time to detection as a function of ΔS for each coloured target/fish during successful trials and ii) number of incorrect pecks during trials. Lines plotted with LOESS (Local Polynomial Regression Fitting) and shaded areas indicate confidence intervals. Additional figures have been provided in the supplementary information suitable for black and white printing.

## Discussion

We have presented a modification of the Ishihara colour “blindness” method for animals, and demonstrate how it can be used to assess colour discrimination thresholds of ateleost. We have provided Matlab code and other resources so that this method may be used with other animals. The method differs fundamentally from most other tests of animal colour vision, which are based on memory, and as such our method is much more similar to the methods used to test human colour thresholds, and the role of colour in visual search. Fish learnt the task quickly, and multiple colours could be tested concurrently due to the fishes’ innate ability to learn the ‘oddity from sample’ methodology, avoiding the need to train them separately to each rewarded colour. Therefore, at a more practical level, many colours can be tested rapidly and concurrently making it possible to investigate colour discrimination throughout colour space in more detail than has hitherto been possible in non-human animals (Green et al. *in prep*.). This method was chosen to resemble natural foraging behavior of the triggerfish i.e. pecking at objects on the substrate, rather than selecting a specific spectral stimulus to receive a food reward, which we expect to be reproducible in other species with comparable foraging ecologies. Indeed, this method could be used to test the visual capabilities of other fish species, birds, and mammals, including standard laboratory model organisms such as rodents and zebrafish. One disadvantage of this method is the significant cost involved in printing large quantities of stimuli using a LaserJet printer and slight variation in printed colours over time. We believe some animals could be trained to instead peck at laminated stimuli or tap at a screen (even a touchscreen) displaying the stimuli to receive a food reward from above, which could be more cost effective and limit the time taken to produce the stimuli. Indeed, our triggerfìsh have since been trained to tap on Ishihara stimuli displayed on an iPad screen placed in an underwater housing.

Our fish were able to perform an oddity colour task and learned to find a dot which differed from the background, regardless of the particular hue. In further experiments, we have found that fish also perform this task when the background is coloured, rather than achromatic grey dots (Green et al, unpublished). Once thought to be a complex cognitive skill restricted to primates, the ability to recognize relationships between stimuli, demonstrated through oddity or sameness/difference learning, has been demonstrated in an increasingly wide range of species. Oddity or sameness/difference tasks have been demonstrated in other fish species, including goldfish (Goldman & Shapiro, 1979) and archerfish (Newport, Wallis, & Siebeck, 2015); several species of birds: such as pigeons (Zentall & Hogan, 1974), crows (Smirnova, Lazareva, & Zorina, 2000) and parrots (Pepperberg, 1987); primates (Katz, Wright, & Bachevalier, 2002)(Bovet & Vauclair, 2001) and bees (Giurfa, Zhang, Jenett, Menzel, & Srinivasan, 2001; Brown & Sayde, 2013).

Our stimuli feature many background dots, which is expected to improve the ease which animals can perform this task, as it has been demonstrated that a large number of distractors improves performance (Zentall et al. 1980), perhaps by increasing recognition of the relationship between distractors, and because they produce a ‘pop out’ effect in which the odd stimuli stands out from the rest. In addition, many vertebrates, for example guppies (Eakley & Houde, 2004) and birds in urban areas (Tryjanowski et al., 2016) have shown an innate preference for novel items, which may attract them to an odd coloured dot. Colour and contrast are crucial cues in animal learning (Newport et al., 2017; Osorio, Jones, & Vorobyev, 1999) and so we anticipate that learning to find an odd colour may be easier than other tasks, such as learning the odd pattern. The use of an oddity from sample method is indeed advantageous because animals learn the task rather than a particular colour and readily generalize to other colours. We envisage that a number of other animals will also be able to perform this behavior with relative ease.

This are mindful that our methodology may be prone to false negatives: if the animals do not respond to the target spot they may still be able to discriminate it from background spots but other factors may influence their decisions. However, our fish were very motivated to perform the task to receive a food reward and made very few mistakes when making a correct choice; therefore, we do not believe that false negatives significantly impacted our results but perhaps should be considered if this method is used with other animals.

To enable accurate calibration of coloured dots, it would be necessary to have spectral sensitivity measurements of cone photoreceptors using microspectrophotometry (MSP) from the animal being tested, in addition to information on spectral filters, including cornea, lens, and oil droplets. Detailed information on how achromatic signals are processed may not be essential if the background dots cover a luminance range that encompasses the brightness of the target colours, as modelled with a range of probable luminance channels (i.e. combined quantum catch of double cone vs quantum catch of long wavelength receptor alone).

Using this methodology, our study demonstrated that discrimination thresholds varied according to the direction of colour space tested, compared to the theoretical prediction (Vorobyev & Osorio 1998). Discrimination thresholds ranged from 0.7 ΔS (Teal) to 2.9 ΔS (Pink). Some of this variation in thresholds may indicate that noise levels, as calculated by the weber fraction and the ratio of different photoreceptor types, were incorrect. Measurements of noise within individual photoreceptors is available for very few species, namely honeybees (Vorobyev 2001), so it would of great value to measure photoreceptor noise in other species. Other factors that may influence thresholds may include co-expression of opsin genes in particular parts of the retina (Dalton, Loew, Cronin, & Carleton, 2014), background colour (adaptation), and/or temporal and spatial effects. Using pairwise tests, discrimination thresholds were measured in our model species *R. aculeatus,* as approximately 2 ΔS (Champ et al. 2016) for blue colours; therefore, both methods appear to give similar results, but this latter experiment only measured thresholds in one area of colour space.

We also recorded time to detection and number of incorrect choices, which can be used to measure the detection of suprathreshold colours and colour saliency. With minor modifications, this methodology could explore grouping of stimuli using achromatic and chromatic cues, or investigate the impact of sensory noise on signal detection by designing background dots so they differ in achromatic and/or chromatic noise. This method could also be used to examine colour categorization and sensory biases.

## Acknowledgements

We thank the staff at Lizard Island Research Station for logistic help on trips to collect the fish and to Cairns Marine Pty Ltd for help in shipping fish to The University of Queensland.

## Authors’ contributions

JAE and NJM conceived the idea; KLC, DCO, NJM, MV and JAE designed the methodology; JAE wrote the MATLAB software to design and produce the stimuli; NFG and AV calibrated the colour stimuli, trained the fish and collected the data; KLC, NFG, and JAE analyzed the data; KLC, NFG, DCO and JAE led the writing of the manuscript. All authors contributed critically to the drafts and gave final approval for publication.

## Data accessibility

Data and Matlab code available from the Dryad Digital Repository http://dx.doi.org/xxxxxx

